# Metabolic modeling unveils potential probiotic roles of *Flavonifractor plautii* in reshaping the Western gut microbiota landscape

**DOI:** 10.1101/2025.04.16.649128

**Authors:** William T. Scott, Enden Dea Nataya, Clara Belzer, Peter J. Schaap

## Abstract

*Flavonifractor plautii*, a prevalent gut commensal, uniquely combines flavonoid degradation with the capacity to produce health-promoting short-chain fatty acids (SCFAs), notably butyrate and propionate. However, its metabolic pathways, ecological roles, and health impacts remain poorly characterized. To explore its probiotic potential and ecological functions, we developed a genome-scale metabolic model (GEM), *iFP655*, using automated reconstruction, deep-learning-based gap-filling, thermodynamic constraints, and transcriptomics. The *iFP655* model substantially improved the predictions of growth rates and SCFA profiles compared to previous models. Simulations identified acetyl-CoA pathways as the preferred route for butyrate production, whereas the energetically costly lysine pathway remained inactive despite robust gene expression. Propionate synthesis occurred primarily via the methylmalonyl-CoA pathway. Community metabolic modeling with representative species of a Western minimal gut microbiota highlighted *F. plautii* ‘s contributions to enhanced SCFA production, especially butyrate, amino acid metabolism, and syntrophic interactions driven by dietary substrates. Our findings indicate that diet-driven syntrophy significantly shapes microbial community structure and function, underscoring the ecological importance of *F. plautii* in gut microbial interactions and highlighting its potential as a probiotic candidate to beneficially modulate gut microbiota through dietary interventions.

## Introduction

The human gastrointestinal tract contains a dense and dynamic community of microorganisms that comprise the gut microbiome, a central component in the maintenance of host health. Dominated by taxa from the phyla Actinobacteria, Bacteroidota, Proteobacteria, Firmicutes, and Verrucomicrobia, these microbial communities engage in essential functions such as maintaining epithelial barrier integrity, modulating immune responses, and transforming otherwise indigestible dietary substrates into bioactive metabolites that influence systemic physiology [40]. Perturbations in the composition and activity of these communities - commonly referred to as dysbiosis - have been implicated in a wide range of chronic diseases, including inflammatory bowel disease, obesity, type 2 diabetes, and neurological conditions [10, 22].

Among the environmental factors shaping gut microbiota composition, diet plays a pivotal role. Western dietary patterns, characterized by high intakes of fat and protein and low levels of dietary fiber, have been associated with gut microbial configurations that exhibit diminished production of short-chain fatty acids (SCFAs) and increased representation of potentially pathogenic bacteria [17, 31]. SCFAs such as acetate, butyrate, and propionate are generated primarily through microbial fermentation of resistant starches and non-digestible fibers, and their physiological relevance spans immune modulation, colonic epithelial maintenance, and energy metabolism [16]. Low levels of SCFAs—particularly butyrate and propionate—are commonly observed in patients with colorectal cancer and inflammatory bowel disease, suggesting that impaired SCFA metabolism may contribute to disease etiology [9].

The growing interest in the therapeutic potential of microbiome modulation has spurred research on SCFA-producing bacteria and their possible deployment as probiotics. Interventions leveraging live microbial strains or prebiotics designed to stimulate their activity have shown promise in elevating SCFA levels and ameliorating disease phenotypes in animal and human models [21, 33]. However, a major barrier to rational strain selection is the incomplete understanding of the metabolic functions and ecological roles of candidate organisms.

A bacterium that has emerged as a promising SCFA producer is *Flavonifractor plautii*, a member of the *Firmicutes* phylum. This organism has been detected in human gut metagenomes and is capable of utilizing substrates such as glucose, maltose, and xylose [54]. Depending on the substrate and growth conditions, *F. plautii* can produce lactate, butyrate, and propionate—metabolites that are of particular interest due to their influence on gut barrier integrity and immune responses. Despite these capabilities, *F. plautii* remains relatively underexplored. Associations with human health, including potential links to obesity and colorectal cancer, have further intensified interest in their functional role within the gut ecosystem [19, 25]. Yet, experimental data on its physiology are limited, and its context-specific interactions with other gut microbes remain unclear.

One major obstacle to elucidating the metabolic capabilities of *F. plautii*—and indeed many gut commensals—is the difficulty of cultivating these organisms in isolation. Many microbes depend on cooperative or competitive interactions within a microbial consortium, rendering monoculture studies insufficient or even infeasible [49]. This constraint has spurred the development and application of computational modeling tools that can infer microbial metabolism *in silico* using genomic information.

Genome-scale metabolic models (GEMs) have become indispensable tools in microbial systems biology. These models systematically represent the entire metabolic network of an organism, including annotated genes, reactions, and metabolites to allow simulation of growth, nutrient utilization, metabolite production, and gene essentiality under defined environmental conditions [13]. Constraint-based modeling approaches such as flux balance analysis (FBA) and thermodynamics-based flux analysis (TFA) enable GEMs to predict feasible metabolic states without requiring detailed kinetic parameters [20]. When integrated with omics data such as transcriptomics, GEMs can be tailored to reflect condition-specific activity, offering mechanistic insights into microbial responses to environmental changes [4, 45].

However, building accurate GEMs remains a non-trivial task, especially for less characterized organisms. Although automated reconstruction tools such as CarveMe and Gapseq facilitate the rapid generation of draft models from genome sequences [29, 56], these drafts often lack key reactions and fail to recapitulate experimentally observed phenotypes. Incompleteness arises from gaps in genome annotation, erroneous assumptions about reaction directionality, or missing transporters. Gap-filling strategies are therefore critical in improving model completeness and predictive power. Traditional gap-filling approaches often require experimental data for calibration, which are unavailable for most gut microbes.

Recent advances in machine learning offer an alternative path to fill gaps in metabolic networks. One such tool, CHESHIRE, applies hypergraph learning to infer missing reactions from the topology of the network itself [8]. By ranking reactions based on confidence and network similarity, CHESHIRE facilitates the judicious addition of reactions to draft models while minimizing the introduction of biologically implausible routes. In tandem, thermodynamic constraints help prevent the emergence of erroneous energy-generating cycles (EGCs) that can artificially inflate biomass predictions [15]. Integrating Gibbs free energy estimates, for example, can help refine reaction directionality and ensure more realistic metabolic simulations [3].

Beyond modeling individual microbes, GEMs have been extended to study microbial consortia, enabling the investigation of species interactions, metabolite cross-feeding, and community-level function [46]. For instance, the Diet-based Minimal Microbiome (DbMM) is a synthetic community comprising ten core gut species selected for their ability to convert Western diet-derived fibers into SCFAs [48]. This system provides a powerful model for exploring how microbial trophic roles shape ecosystem function. In such communities, *F. plautii* has been suggested to play a role in the downstream metabolism of amino acids and fermentation by-products, particularly under carbohydrate limitation conditions [49]. Simulations involving DbMM subcommunities can shed light on syntrophic relationships and help identify keystone interactions within gut ecosystems.

Despite these methodological advancements, a curated and context-specific GEM for *F. plautii* has not yet been developed. Publicly available models, such as AGORA v1.03 entry, often lack manual curation and do not reliably predict metabolic fluxes, especially under gut-relevant conditions [55]. This gap impedes efforts to mechanistically characterize *F. plautii*’s contributions to gut microbiota structure and function of the intestinal microbiota or to evaluate its suitability as a probiotic candidate.

In this study, we reconstructed a high-quality genome-scale metabolic model of *Flavonifractor plautii*, designated *iFP655*. Using both CarveMe and Gapseq, we generated and compared draft models that were subsequently refined through manual curation, integration of thermodynamic constraints, and topology-guided gap filling using CHESHIRE. To contextualize the model under gut-relevant conditions, transcriptomic data from a synthetic Diet-based Minimal Microbiome (DbMM) community were incorporated via the GIMME algorithm. The model’s ability to simulate SCFA production and amino acid utilization was validated through *in silico* experiments using parsimonious flux balance analysis and flux variability analysis under various media conditions. Furthermore, we reconstructed the GEMs for other DbMM members and applied them in a three-species community simulation with *Bacteroides ovatus* and *Coprococcus catus* to examine potential syntrophic interactions. The resulting model offers a mechanistic framework for understanding *F. plautii*’s metabolic potential and its ecological roles in SCFA production and cross-feeding of nutrients within the gut microbiome.

## Results

### Flavonifractor plautii GEM reconstruction

The *F. plautii* model reconstructed by CarveMe has a slightly higher number of genes and reactions compared to the Gapseq model. However, the Gapseq model predicted that more metabolites would be present in the draft model (1,445 metabolites), while CarveMe predicted only 1,171 metabolites. The properties comparison of both *F. plautii* draft models is summarized in Table 1. Both CarveMe and Gapseq draft models were further curated through manual curation and an automated gap-filling process using CHESHIRE, resulting in 10 different models for *F. plautii* (i.e., 2 manually curated models and 8 gap-filled models).

**Table 1.**
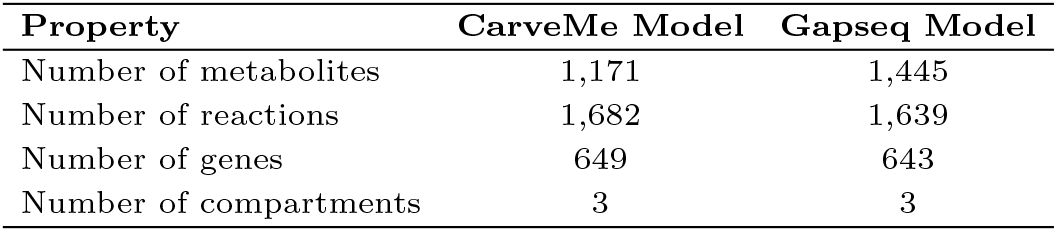
*Flavonifractor plautii* draft model properties comparison.

| Property | CarveMe Model | Gapseq Model |
| --- | --- | --- |
| Number of metabolites | 1,171 | 1,445 |
| Number of reactions | 1,682 | 1,639 |
| Number of genes | 649 | 643 |
| Number of compartments | 3 | 3 |

To evaluate the models, we compared *F. plautii* predicted growth rates in three different media with experimental data [1, 48, 55]. Parsimonious flux balance analysis (pFBA), with biomass reaction set as the objective function, revealed that both the manually curated CarveMe and Gapseq models underestimated growth rates in AF and M2 media, while overestimating the growth rate in DbMM medium (Figure 1).

**Fig. 1.**
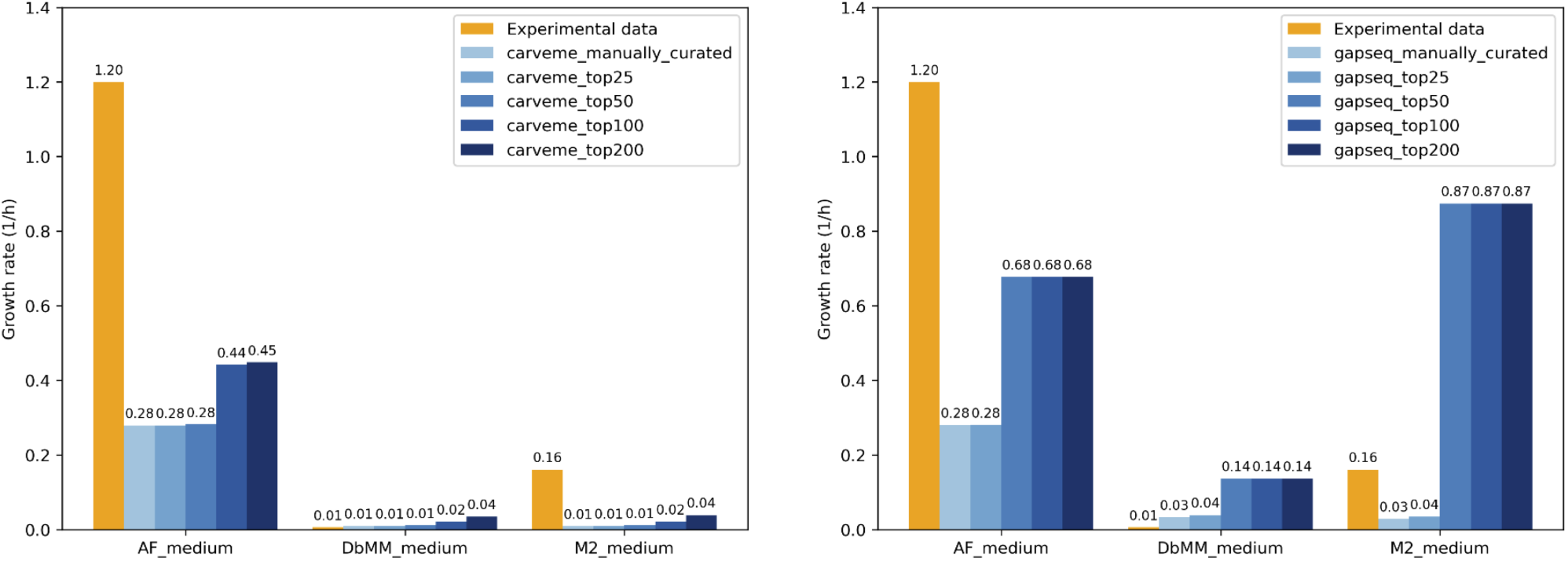
Comparison of predicted growth rates for *Flavonifractor plautii* models reconstructed using CarveMe (left) and Gapseq (right) against experimental measurements in AF, DbMM, and M2 media. The baseline draft models were manually curated and subsequently refined using the CHESHIRE tool, a deep learning-based gap-filling approach. Each bar represents a model variant, where `*_manually_curated` refers to the curated draft prior to gap filling, and `*_top25`, `*_top50`, `*_top100`, and `*_top200` correspond to the same model augmented with the top 25, 50, 100, or 200 CHESHIRE-predicted reactions, respectively. The orange bars represent experimentally observed growth rates.

This suggests that the gap-filling process during model reconstruction was optimized primarily for the medium used during the gap-filling (i.e. DbMM medium), potentially leading to discrepancies in other conditions. Furthermore, pFBA results showed that gap-filled models resulted in higher predicted growth rates. Notably, the Gapseq models exhibited a sharp increase in growth rate upon the addition of the top 50 reactions. However, no further changes in growth rate were observed in models with more added reactions, indicating that the first 50 gap-fill reactions were crucial for biomass synthesis, while the remaining 150 may participate in other metabolic pathways. The list of top 200 candidate reactions for both CarveMe and Gapseq models is provided in Supplementary Table S2.

Simulations also showed that adding too many reactions (i.e., top 100 and 200) led to inflated growth rate predictions for both the CarveMe and Gapseq models. In contrast, models with fewer reactions (top 25 and 50) yielded more realistic predictions, especially for the CarveMe-reconstructed models. The `carveme_top50` model yielded the closest growth rate prediction in DbMM medium, with a predicted value of 0.0099 h^-1^, compared to the experimental value of 0.0059 h^-1^.

In addition to comparing growth rates, we also evaluated the main short-chain fatty acid (SCFA) secretion profiles predicted by the models against experimental data [48]. All CarveMe-based models failed to predict the secretion of acetate. Although butyrate and propionate secretion were qualitatively predicted, the corresponding flux values deviated from the experimental measurements. In particular, the addition of the top 200 gap-fill reactions to the CarveMe model further exacerbated the deviations, leading to overestimated fluxes for both butyrate and propionate (Figure 2).

**Fig. 2.**
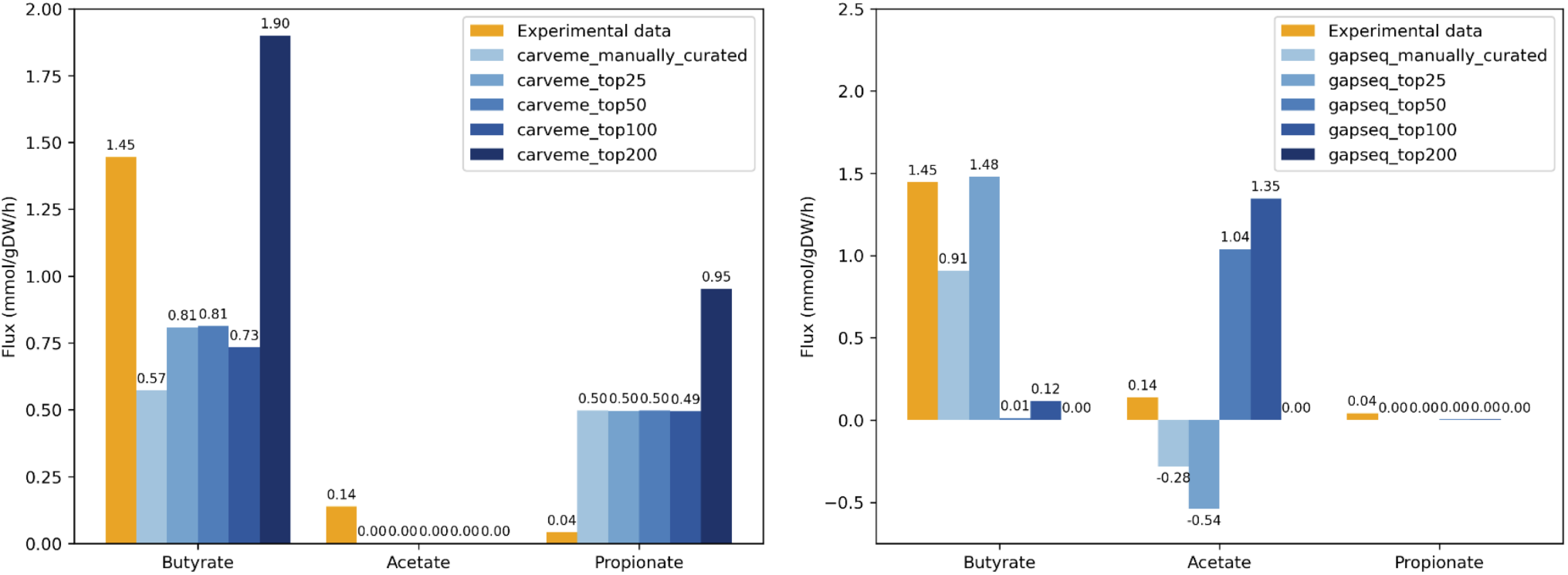
Comparison of predicted metabolite fluxes (mmol/gDW/h) from *Flavonifractor plautii* genome-scale metabolic models reconstructed using CarveMe (left) and Gapseq (right) against experimental data for butyrate, acetate, and propionate. Each model was further refined by gap-filling with CHESHIRE, an automated gap-filling algorithm that integrates machine learning with biochemical context. Bars labeled `manually_curated` represent models refined through manual curation, while `top25`, `top50`, `top100`, and `top200` denote automated CHESHIRE gap-filling using the top 25, 50, 100, and 200 ranked candidate reactions, respectively, based on model-guided scoring.

The `gapseq_manually_curated` model failed to predict propionate production. Furthermore, it predicted consumption of acetate instead of secretion. Adding gap-fill reactions appeared to shift the flux distribution from butyrate production to acetate, as shown in the SCFA secretion profiles of the `gapseq_top50` model. Meanwhile, propionate secretion was only observed in the `gapseq_top50` and `gapseq_top100` models, and at very low flux levels (i.e., 0.003574 mmol/gDW/h for both models) (Figure 2).

These results indicate that none of the reconstructed models fully recapitulates the experimental data. However, compared to publicly available models—such as the *F. plautii* ATCC 29863 AGORA v1.03 model and the *F. plautii* YL31 model from a previous study [55]—our models showed improved accuracy in predicting both growth rate and SCFA production in the DbMM medium. The AGORA model failed to predict any growth due to the absence of essential metabolites in the defined medium, while the YL31 model overestimated the growth rate nearly eightfold (0.0477 h^-1^). Additionally, the YL31 model predicted acetate as the primary SCFA produced (acetate exchange flux of 1.109 mmol/gDW/h), underestimated butyrate production (0.37 mmol/gDW/h), and did not predict propionate secretion at all (Table 2).

**Table 2.**
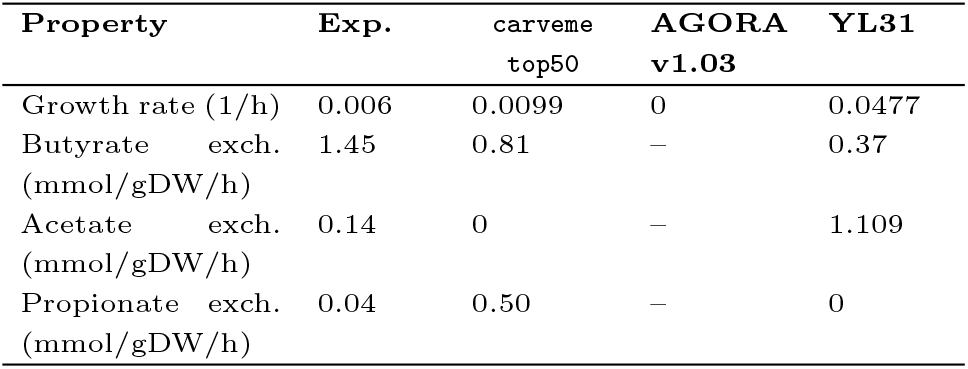
Comparison of predicted growth rates and metabolite fluxes between the carveme_top50 model and published models of *F. plautii*.

| Property | Exp. | carveme<br>top50 | AGORA<br>v1.03 | YL31 |
| --- | --- | --- | --- | --- |
| Growth rate (1/h) | 0.006 | 0.0099 | 0 | 0.0477 |
| Butyrate exch.<br>(mmol/gDW/h) | 1.45 | 0.81 | – | 0.37 |
| Acetate exch.<br>(mmol/gDW/h) | 0.14 | 0 | – | 1.109 |
| Propionate exch.<br>(mmol/gDW/h) | 0.04 | 0.50 | – | 0 |

These discrepancies highlight that both the growth rate and SCFA secretion profiles predicted by the AGORA and YL31 models were less accurate than those produced by our models. Consequently, the `carveme_top50` model was selected as the optimal model in this study, as it produced growth rate and butyrate flux predictions, particularly in DbMM medium, that were closest to experimental values. This model was used for subsequent analyses.

### Essential gene identification and subsystem distribution in the **F. plautii** GEM

The metabolic subsystem distribution of the *F. plautii* model is shown in Figure 3 (details in Supplementary Table S1). Transport reactions represented the largest metabolic subsystem in the model, accounting for 439 reactions, or approximately 25.24% of the total reactions. This was followed by amino acid metabolism (approximately 16.50%) and carbohydrate metabolism (approximately 12.24%).

**Fig. 3.**
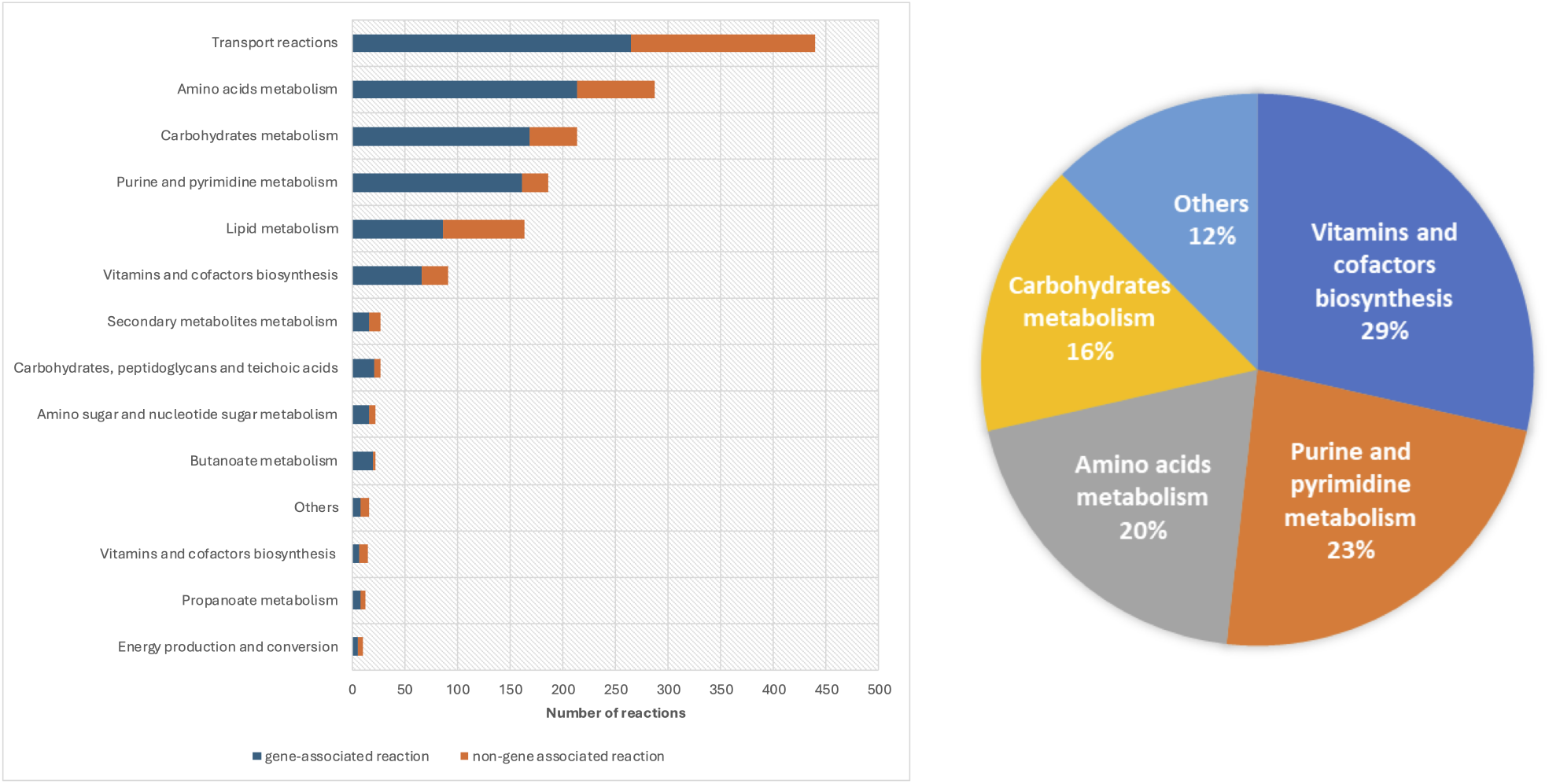
(Left) Metabolic subsystem distribution of the *Flavonifractor plautii* model, including gene- and non-gene-associated reactions. (Right) Distribution of metabolic pathways associated with the 56 essential genes identified through *in silico* single-gene deletion analysis under DbMM medium conditions.

The majority of non-gene-associated reactions were found within the transport subsystem. This is expected as many transport processes, such as the movement of metabolites between the periplasm and cytosol, may occur spontaneously and thus lack direct gene associations. Lipid metabolism, amino acid metabolism, and carbohydrate metabolism also contributed substantially to the pool of non-gene-associated reactions, accounting for 78, 74, and 45 internal reactions, respectively.

Gene essentiality analysis identified 56 genes as essential for the growth of *F. plautii* under DbMM medium conditions (see Supplementary Table S3). The metabolic subsystems associated with these essential genes are illustrated in Figure 3.

### Transcriptomics data integration improved butyrate prediction

The initial transcriptomics dataset for *F. plautii* from a previous study included 5,485 genes expressed [49]. After filtering to retain only the top 30th percentile, 1,646 genes remained, with expression levels ranging from 4.36 to 2,345 counts per million (CPM) (Supplementary Table 10 in [49]). Reactions associated with these genes were assumed to be active, as indicated by their relatively high expression levels. Integrating transcriptomics data into the model resulted in the knockout of 351 reactions (Supplementary Table S3).

This adjustment slightly increased the predicted growth rate to 0.0105 h^-1^. In particular, the prediction of butyrate exchange flux improved: the transcriptomics-integrated model estimated a flux of 1.29 mmol/gDW/h, closely aligning with the experimental value of 1.45 mmol/gDW/h, and improving upon the previous model’s prediction of 0.81 mmol/gDW/h. This enhancement demonstrates the importance of transcriptomics data in guiding fluxes toward biologically relevant pathways. By incorporating gene expression, the model more accurately reflected the metabolic activity of the organism, leading to better prediction of key metabolites such as butyrate. This transcriptomics-integrated model was used for all subsequent analyzes.

### Acetyl-CoA and lysine pathways are primary routes for butyrate production in **F. plautii**

Two butyrate biosynthesis pathways were included in the model: the acetyl-CoA pathway and the 4-aminobutanoate /succinate pathway. Steady-state simulation in DbMM medium indicated that the acetyl-CoA pathway was the primary route. The terminal step was catalyzed by butyryl-CoA: acetate CoA transferase (BUTCT2) rather than butyrate kinase (BUTKr). The BUTCT2 reaction facilitates the transfer of CoA between butyryl-CoA and acetoacetate, producing butyrate and acetoacetyl-CoA. The genes associated with BUTCT2, `WP_007495948_1` and `WP_007495949_1`, were within the top 30^th^ percentile expressed genes, with 76.35 and 72.19 CPM, respectively. These genes encode CoA-transferase subunits A and B. In contrast, the BUTKr reaction was knocked out due to low gene expression (2.13 CPM). Furthermore, four reactions in the 4-aminobutanoate pathway—`SSCOARy`, `SSAH`, `A4HBCT`, and `4HBCOAH`—were also removed due to low expression levels.

The lysine utilization pathway was not initially present in the CarveMe-reconstructed model, despite the fact that the relevant genes are encoded in the genome. This probably stems from the absence of the pathway in the universal CarveMe model, which was based on the BiGG database circa 2018 [29]. The complete lysine-to-butyrate pathway was only later introduced in the *Clostridioides difficile* iCN900 model published in 2020 [35].

To address this gap, five reactions from the lysine pathway were manually added to the model along with the corresponding gene-protein-reaction (GPR) rules. These are listed in Table 3. The final model, named *iFP655*, consisted of 1,198 metabolites, 1,739 reactions, and 655 genes.

**Table 3.** Reactions from the lysine utilization pathway incorporated into the *F. plautii* model.

| Reaction ID | Reaction Name | GPR |
| --- | --- | --- |
| LYSAM | Lysine 2,3-aminomutase | WP_009258627.1 |
| DH36M | Lysine 5,6-aminomutase | WP_009258626.1 and WP_007493054.1 |
| DH35O | 3,5-diaminohexanoate dehydrogenase | WP_007493057.1 |
| A53C | 3-keto-5-aminohexanoate cleavage | WP_007493059.1 |
| AB3CL | 3-aminobutyryl-CoA ammonia lyase | WP_009259558.1 or WP_007488193.1 |

Transcriptomics data confirmed high expression of all genes involved in the lysine pathway, with transcript levels ranging from 83.95 to 319.46 counts per million (CPM). However, parsimonious flux balance analysis (pFBA) simulations showed that integrating this pathway did not alter the flux distribution of the model; both the predicted growth rate and the butyrate production remained unchanged. All fluxes through the lysine pathway were zero, indicating that the acetyl-CoA pathway remained the preferred route for butyrate biosynthesis. This may be due to the dual role of lysine as a biomass precursor and its higher energetic cost: the lysine pathway requires ATP (for example, through the LYSabc transport reaction), whereas the acetyl-CoA pathway does not. In the nutrient-limited DbMM medium, the model appears to favor the more energy-efficient acetyl-CoA route.

Flux variability analysis (FVA) suggested a possible upper bound activity for the lysine pathway, with a maximum flux of 0.05 mmol/gDW/h. The butyrate exchange reaction exhibited a flux range of 0.7376 to 1.8211 mmol/gDW/h (Supplementary Table 7). The corresponding flux distribution and gene expression levels in the butyrate and propionate synthesis pathways are presented both as a simplified metabolic overview (Figure 4) and as a detailed map of annotated fluxes and transcriptomic data (Figure 5).

**Fig. 4.**
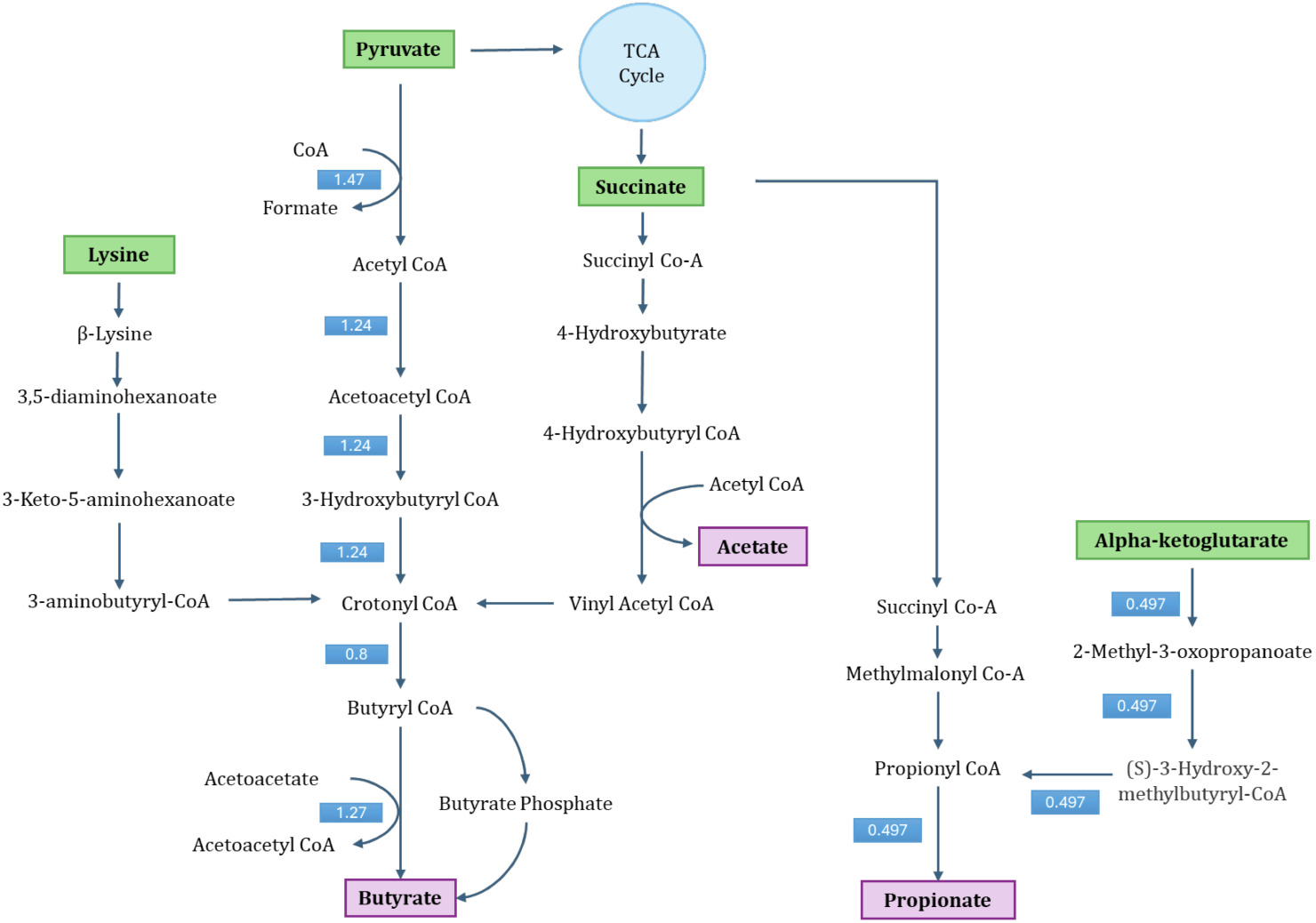
Simplified flux distribution map of butyrate and propionate synthesis pathways in *Flavonifractor plautii*. Flux values (in mmol/gDW/h) are indicated along key reactions involved in short-chain fatty acid biosynthesis.

**Fig. 5.**
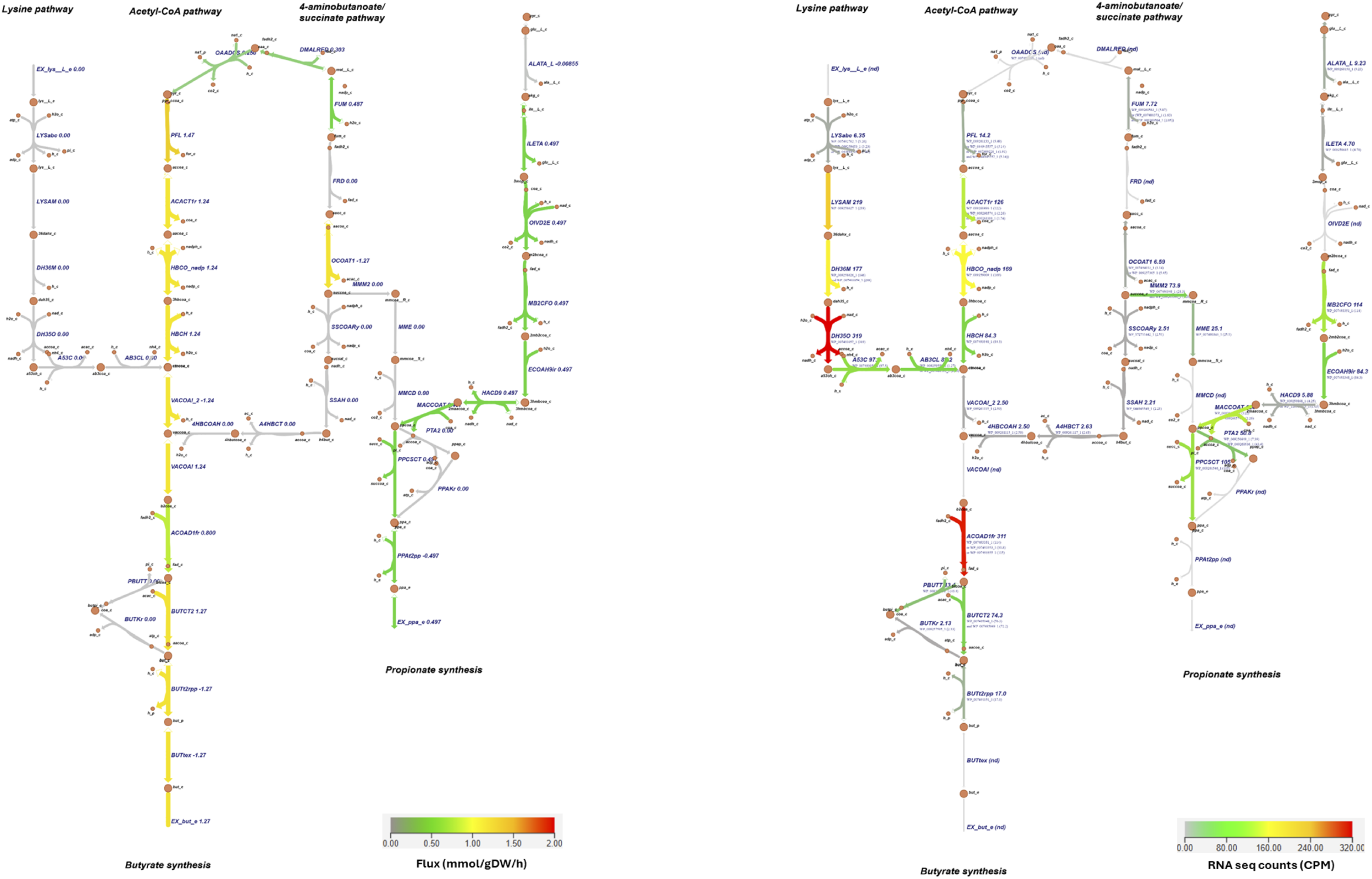
Flux distribution (left) and gene expression (right) maps of butyrate and propionate synthesis pathways in *Flavonifractor plautii*. Each arrow represents a reaction, with color intensity proportional to either predicted flux (mmol/gDW/h, left) or transcript expression (CPM, right).

Comparison of predicted pFBA fluxes and transcriptomics data under DbMM conditions revealed strong alignment for most pathways, except the lysine pathway. The acetyl-CoA pathway showed high flux and high gene expression, particularly for key enzymes such as acetyl-CoA acetyltransferase (ACACT1r), 3-hydroxybutyryl-CoA dehydrogenase (HBCO nadp), and short-chain enoyl-CoA hydratase (HBCH). In contrast, the 4-aminobutanoate/succinate pathway showed no flux and low gene expression, indicating its inactivity. Despite strong transcriptomic support for lysine pathway genes—including `LYSAM`, `DH36M`, `DH35O`, `A53C`, and `AB3CL`—the pathway was not used in simulations, suggesting that under growth-limiting conditions, lysine is preferentially channeled into biomass formation rather than fermentation.

Transcriptomics data also suggested potential activity of the methylmalonyl-CoA pathway for propionate production. In this route, succinate is first converted to R-methylmalonyl-CoA by methylmalonyl-CoA mutase (MMM2), then rearranged to S-methylmalonyl-CoA by epimerase (MME), and finally decarboxylated to propionyl-CoA by methylmalonyl-CoA decarboxylase (MMCD). Propionyl-CoA is subsequently converted to propionate via propionyl-CoA:succinate CoA-transferase (PPCST). However, predicted fluxes through MME, MMM2, and MMCD were zero, indicating a discrepancy between gene expression and metabolic activity.

An alternative route for propionate production supported by both gene expression and flux data involved alpha-ketoglutarate (AKG) metabolism via branched-chain amino acid (BCAA) pathways. Although not a direct route, AKG can be converted to propionyl-CoA, a precursor to monomethyl BCAAs, which in turn feeds into the PPCST reaction for propionate synthesis [27]. This result highlights a potential link between amino acid metabolism and short-chain fatty acid (SCFA) biosynthesis.

The model did not simulate acetate production, probably because of the limited energetic benefit of acetate secretion under DbMM conditions. Two key reactions that contribute to the formation of acetate, 4-hydroxybutanoyl-CoA dehydratase (4HBCOAH) and acetyl-CoA: butyrate-CoA transferase (BUTCT were both inactive in the simulations. In addition, the 4HBCOAH reaction was removed from the model due to low gene expression. These results suggest that the model’s underestimation of acetate production may stem from an incomplete representation of alternative acetate biosynthesis routes.

### The reconstruction pipeline lacks robustness for all community member models

The same pipeline used to generate *iFP655* was applied to reconstruct genome-scale metabolic models for the nine remaining species in the DbMM community. The general reconstruction workflow is shown in Figure 8. Initially, this pipeline failed to predict growth for three species: *E. siraeum, S. variabile*, and *B. ovatus*. These models contained internal oxygen-dependent reactions that were essential for growth (see supplementary material, Table S4).

To resolve this, we restored the original upper and lower bounds of these oxygen-dependent reactions to their pre-curation values while maintaining the oxygen exchange reaction (`EX_o2_e`) as blocked. This adjustment enabled non-zero growth in all models (Figure 6).

**Fig. 6.**
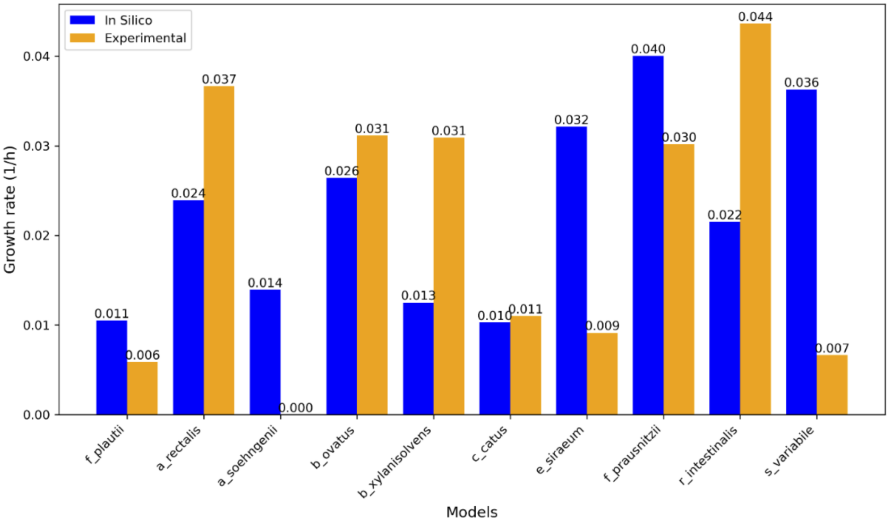
Comparison of growth rates predicted by genome-scale metabolic models (GEMs) with experimentally measured growth rates.

Steady-state simulations using pFBA showed that although the pipeline could yield viable models for all members of the DbMM community, the predicted growth rates varied widely compared to the experimental data. The most accurate prediction was achieved by the *C. catus* model, which had a predicted growth rate of 0.0103 h^-1^, closely matching the experimentally observed value of 0.011 h^-1^. The *B. ovatus* model also yielded a reasonably accurate prediction, albeit slightly underestimating the actual growth rate.

In contrast, several models showed substantial deviations. Most notably, the *A. soehngenii* model predicted a growth rate of 0.014 h^-1^, despite no measurable growth was observed in the corresponding experimental data [48]. These findings suggest that the current reconstruction pipeline lacks sufficient robustness to reliably produce accurate metabolic models for all members of the DbMM community. More manual curation and refinement will be required to improve model accuracy and predictive power.

### Community modeling reveals potential cross-feeding interactions between **F. plautii** and other gut bacteria

A three-species community consisting of *F. plautii, B. ovatus*, and *C. catus* simulated under DbMM medium conditions was predicted to achieve a collective growth rate of 0.231 h^-1^. In the FBA simulation, where species abundances were fixed as input parameters, *B. ovatus* exhibited the highest individual growth rate (0.136 h^-1^), followed by *C. catus* (0.060 h^-1^) and *F. plautii* (0.035 h^-1^). All models predicted higher growth rates in community settings compared to monoculture simulations, suggesting possible cross-feeding interactions that create a more favorable metabolic environment for growth.

The total production of propionate by the community was predicted to be 2.96 mmol/gDW/h, with *B. ovatus* as the primary contributor (1.77 mmol/gDW/h), followed by *F. plautii* (1.36 mmol/gDW/h). In contrast, *C. catus* was predicted to consume propionate at a rate of 0.17 mmol/gDW/h. Acetate was predicted to be produced by both *B. ovatus* (8.25 mmol/gDW/h) and *C. catus* (3.63 mmol/gDW/h). *F. plautii* was the only predicted butyrate producer, with a flux of 1.76 mmol/gDW/h, leading to butyrate accumulation in the community medium. However, no butyrate cross-feeding was observed. Although *C. catus* is a known butyrate producer [36], it did not produce butyrate in the community model. For detailed FBA results for the three-species community, see Supplementary Table 8.

FVA results revealed several potential cross-feeding interactions, as illustrated in Figure 7 (see also Supplementary Table 9). *B. ovatus* was predicted to be the sole consumer of cellobiose. Both *B. ovatus* and *C. catus* consumed xylan, suggesting a shared ecological niche for this polysaccharide. Inulin, starch, and pectin were not utilized in the simulation, likely due to their absence in the individual models and thus also in the combined community model.

**Fig. 7.**
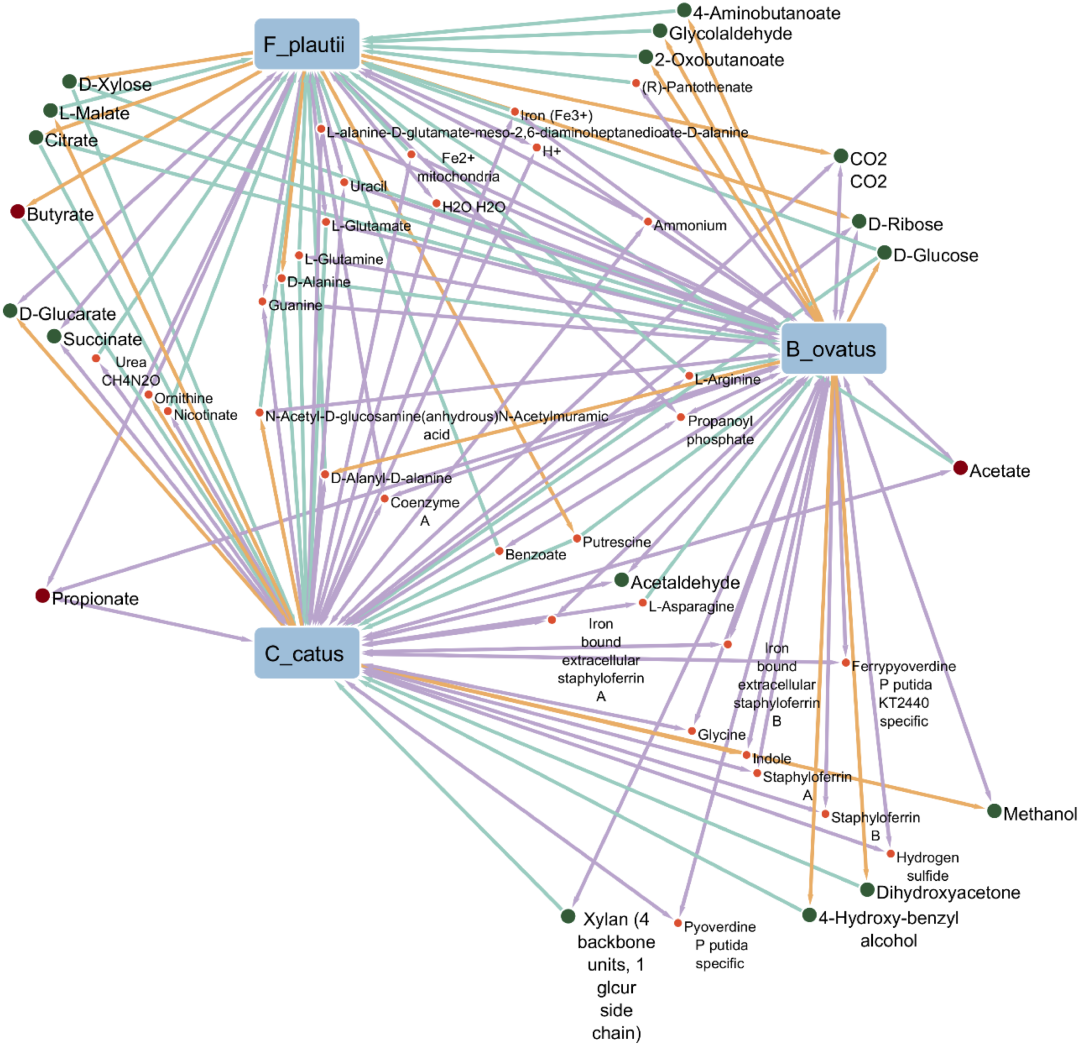
Visualization of the three-species DbMM community model and its predicted cross-feeding interactions based on FVA results. Orange lines indicate production, green lines indicate consumption, and purple lines indicate both production and consumption. Major SCFAs (i.e., butyrate, acetate, and propionate) are marked as red dots, while potential carbon sources are marked as green dots.

**Fig. 8.**
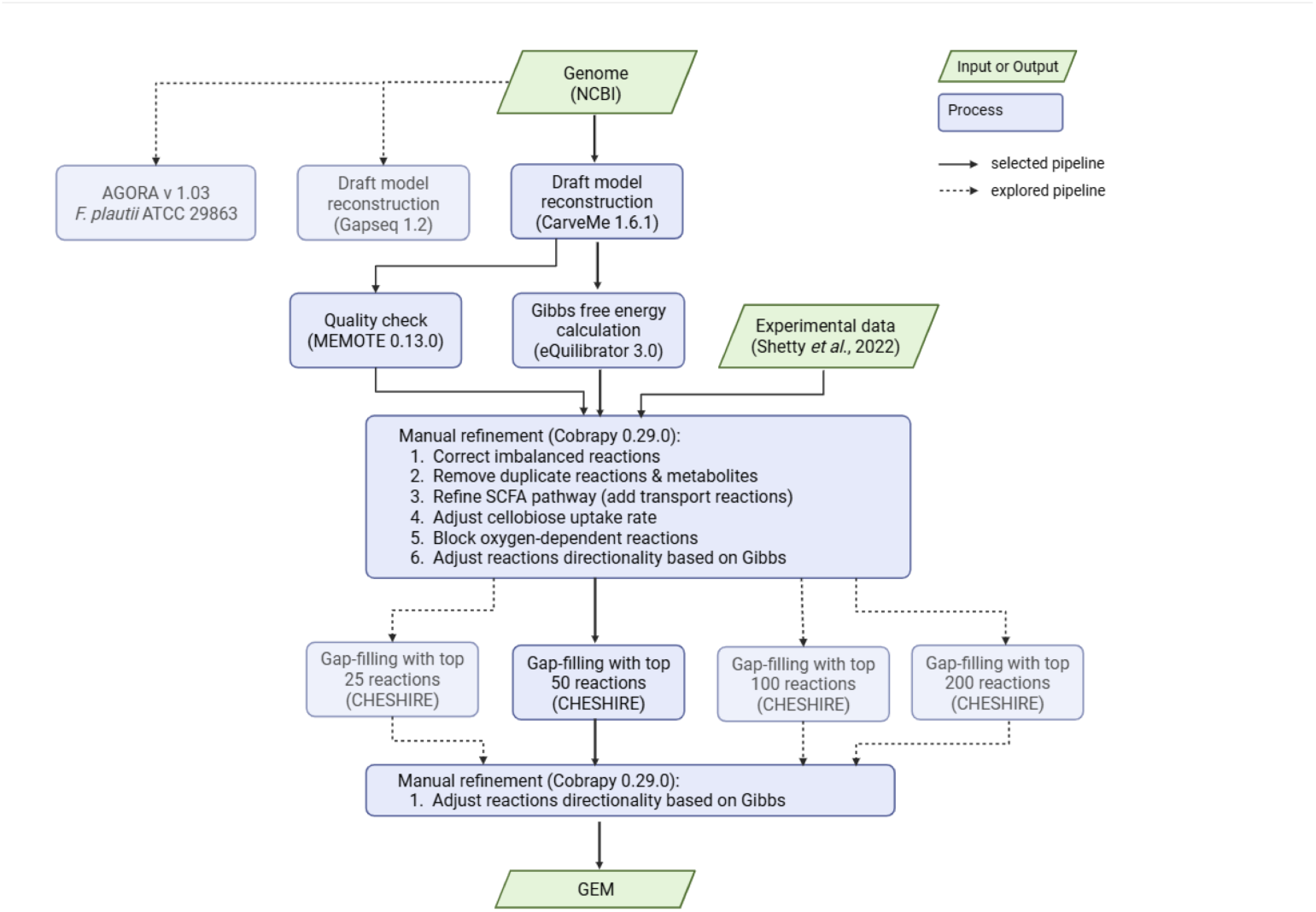
Workflow outlining the genome-scale metabolic model reconstruction pipeline used in this study. The selected pipeline integrates draft model generation (CarveMe or Gapseq), manual curation, thermodynamic constraints, and CHESHIRE-based gap-filling to produce refined GEMs. Alternative pathways explored are indicated with dashed arrows. This framework was applied to reconstruct and evaluate models of *F. plautii* and other Diet-based Minimal Microbiome (DbMM) members.

*B. ovatus* functioned as a primary degrader of complex polysaccharides such as xylan and cellobiose, converting them into simpler sugars like D-ribose and D-glucose, which can serve as public goods for the other species. It also produced acetaldehyde at a maximum rate of 11.14 mmol/gDW/h, which was completely consumed by *C. catus* at the same rate. In addition, *C. catus* secreted L-malate that could be utilized by both *F. plautii* and *B. ovatus*, while succinate was both produced and consumed by *C. catus* and *F. plautii*. Other metabolites, such as D-glucarate, D-xylose, and methanol, were also predicted to participate in low-level cross-feeding interactions, although their ecological significance may be limited by low fluxes.

The three species were predicted to produce and consume propionate, with fluxes ranging from -2.07 to 6.63 mmol/gDW/h. The net accumulation of propionate in the community reached a maximum of 7.68 mmol/gDW/h, suggesting dynamic exchange and partial environmental retention. Acetate was the most abundantly produced metabolite, with a community-level maximum flux of 31.27 mmol/gDW/h. The major contributors were *C. catus* and *B. ovatus*, while *F. plautii* was predicted to consume acetate at a maximum rate of 3.13 mmol/gDW/h, implying its role as a secondary acetate user in the community.

Butyrate production was exclusively attributed to *F. plautii*, which had a maximum predicted flux of 5.91 mmol/gDW/h, reinforcing its ecological role as the primary butyrate producer. Interestingly, *C. catus* was predicted to consume butyrate at a rate of 2.23 mmol/gDW/h, which contrasts with literature reports identifying it as a butyrate producer [36]. Additionally, *B. ovatus* produced 4-aminobutanoate at a rate of 0.99 mmol/gDW/h, which was fully consumed by *F. plautii*. Notably, 4-aminobutanoate is one of the four primary substrates known to support butyrate synthesis. This predicted exchange suggests a potential cross-feeding mechanism that supports butyrate production in the community.

## Discussion

This study presents a genome-scale metabolic model (GEM) for the butyrate-producing gut bacterium *Flavonifractor plautii*, reconstructed using a semi-automated pipeline incorporating automated tools, deep learning-based gap filling, thermodynamic constraints, and transcriptomic data. The resulting model, *iFP655*, allows mechanistic insight into the metabolic functions of the organism and its ecological roles within microbial communities. Our findings also highlight broader considerations for GEM reconstruction of under-characterized gut microbes.

### Curation remains critical for accurate GEM reconstruction

Despite the growing use of automated pipelines, reconstructing GEMs with robust quantitative predictive capacity remains challenging, particularly for under-characterized microbes [18]. In this study, a CarveMe-generated draft refined through CHESHIRE-based gap filling (the top 50 reactions) and manual curation yielded *iFP655*, which accurately predicted growth rate and butyrate production in DbMM medium. Compared to publicly available models (AGORA v1.03 and YL31), *iFP655* demonstrated enhanced predictive performance, with key reactions supported by gene-protein-reaction (GPR) associations.

However, applying the same reconstruction pipeline to other species in the DbMM community revealed significant variability in predicted growth phenotypes. This underscores the limitations of generic pipelines and the importance of targeted curation to improve model accuracy. Several critical aspects were identified to enhance model refinement: precise medium definition, application of thermodynamic constraints, integration of transcriptomics, and careful evaluation of gap-filled reactions.

#### Model behavior influenced by media and thermodynamics

Defining the composition of the medium is essential because it directly constrains the exchange fluxes. In the absence of measured uptake rates, compound concentrations were used as proxies, consistent with previous studies [30]. However, this approach may overestimate substrate availability. Furthermore, the use of complex media, rather than defined formulations, introduces uncertainty due to the poorly characterized nutrient composition. Previous work has shown that such conditions can bias the filling of the gap towards false positive or false negative reactions [6].

Imposing thermodynamic constraints by assigning directionality based on the Gibbs free energy change improved the physiological realism of flux predictions. Reactions with Δ_*r*_*G*° between ±30 kJ/mol were treated as reversible, a convention that aligned with previous modeling studies [5, 20]. This intervention reduced the occurrence of energy-generating cycles (EGCs), a common artifact in draft or uncurated GEMs [15]. Although our estimations were based on standard conditions, future implementations using Thermodynamics-based Flux Analysis (TFA) could further enhance the predictive power [47].

#### Transcriptomics guides model accuracy

Integrating transcriptomics data allowed us to contextualize model predictions under specific growth conditions. Using a 30th percentile expression threshold, we deactivated reactions associated with lowly expressed genes - temporarily turning them off in the simulation without removing them from the model structure - to refine predictions of SCFA fluxes. This approach also revealed inconsistencies in initial reaction assignments, such as the unsupported inclusion of PPAKr in propionate synthesis, which were corrected by enforcing GPR-consistent alternatives. Although our thresholding approach reflects common practice [45], future models may benefit from probabilistic weighting methods, such as RIPTiDe [23].

#### Gap-filling must balance completeness and plausibility

Although CHESHIRE enabled gap filling significantly improved model completeness, it also introduced risks. Adding too many reactions (e.g., top 100–200) resulted in biologically implausible flux distributions, while too few (e.g., top 25) yielded incomplete networks. We found that a selective addition of 50 reactions provided the best trade-off. Notably, some candidate reactions lacked mass balance, as reported by MEMOTE. This reinforces the need to validate gap-filled content against genomic or functional annotations [8].

### Key metabolic capabilities captured in **iFP655**

#### Metabolic versatility and amino acid utilization

*iFP655* successfully recapitulates three canonical butyrate synthesis pathways, acetyl-CoA, lysine, and 4-aminobutanoate / sucinate, each with substantial support from GPR [51]. In addition, it includes the methylmalonyl-CoA pathway for propionate biosynthesis and suggests a connection between BCAA metabolism and SCFA production. These capabilities position *F. plautii* as a metabolically versatile member of the gut microbiome, contributing to both the butyrate and propionate pools [32].

Flux simulations revealed that in DbMM medium, which lacks readily fermentable carbohydrates, *F. plautii* preferentially consumed amino acids such as L-aspartate, L-glutamate, L-isoleucine, L-threonine, and L-valine. These findings align with experimental observations of widespread amino acid catabolism by gut microbes [43] and previous *in silico* predictions showing active degradation pathways for isoleucine, leucine, and tryptophan in *F. plautii* [49]. This highlights the adaptation of the organism to protein-rich, carbohydrate-poor niches and supports its ecological role in nitrogen and energy cycling.

#### Condition-driven lysine-to-butyrate flux

Our simulations indicate minimal usage of the lysine-to-butyrate pathway under the evaluated conditions (*DbMM medium*), primarily due to the presence of energetically favorable alternatives such as the acetyl-CoA pathway. However, under conditions of severe carbohydrate limitation [7], lysine derivatives such as fructoselysine might become viable substrates for butyrate synthesis. Future modeling efforts should investigate nutrient-limiting conditions inspired by diets enriched in advanced glycation end-products (AGEs) [53] or distinct dietary regimens, such as infant formulas versus breast milk [52], to uncover conditions that favor lysine utilization.

Transcriptome-informed refinements further improved model behavior. In particular, disabling the PPAKr reaction (without gene evidence) and enforcing flux through the gene-supported PPCST reaction corrected for inconsistencies in propionate pathway predictions. Future models may benefit from additional omics layers, including proteomics or metabolomics, to further validate network activity under variable conditions.

Despite transcriptomic support for the lysine-to-butyrate pathway, our model simulations did not predict significant flux through this pathway, possibly due to its higher energetic cost compared to alternatives such as the acetyl-CoA pathway. This highlights the importance of combining metabolic modeling with carefully controlled experimental conditions designed specifically to induce targeted pathways.

Finally, *in silico* single-gene deletion analysis identified 56 genes as essential for growth under DbMM medium. Most were involved in the biosynthesis of biomass precursors. Comparative analysis against the ePath database revealed that 45 of these genes were also predicted to be essential *in silico* [2, 24], supporting the accuracy of our predictions and providing a foundation for future functional studies.

### F. plautii contributes to community-level metabolic cooperation

To assess its ecological role, we simulated a three-member community of *F. plautii, B. ovatus*, and *C. catus* under DbMM medium. These species represent trophic guilds 3, 1, and 2, respectively [49]. Our results show that metabolic roles were preserved within the community: *B. ovatus* degraded complex polysaccharides, *C. catus* utilized simple sugars, and *F. plautii* consumed malate, glucose, and acetate. These interactions align with previous classifications of trophic strategies and suggest that *F. plautii* acts as a fermentation by-product utilizer [49, 41].

*F. plautii* remained the sole butyrate producer in the simulated community, with a predicted flux of 1.76 mmol/gDW/h —higher than in monoculture. FVA results indicated a potential maximum flux of 5.91 mmol/gDW/h. The production of propionate in the community was also significantly elevated, reflecting cooperative metabolic enhancement and in agreement with previous studies suggesting that propionate synthesis is more favorable in community settings [41]

These results also suggest metabolite-level cross-feeding: *C. catus* was predicted to consume butyrate, while *B. ovatus* secreted acetaldehyde, which was detoxified by *C. catus* via aldehyde dehydrogenase. This interaction may mitigate the harmful accumulation of acetaldehyde in the gut, a metabolite implicated in gastrointestinal carcinogenesis [34]. Such syntrophic dynamics highlight the potential of SCFA producers like *F. plautii* to support beneficial community configurations.

Our community simulations highlight significant diet-driven syntrophic interactions. Specifically, the metabolism of *F. plautii* could shift dramatically under nutrient-limiting scenarios, altering the ecological landscape through metabolic cross-feeding. Further studies incorporating varied dietary contexts, especially those that mimic Western dietary patterns with altered carbohydrate-to-protein ratios, may elucidate how lysine metabolism could contribute to community stability and SCFA production.

### SCFA production and therapeutic relevance

Short-chain fatty acids, particularly butyrate, are critical for gut health. Butyrate promotes mucosal integrity, suppresses inflammation, and selectively shapes the composition of the microbiota by supporting beneficial taxa and suppressing opportunistic pathogens [44, 50]. In our simulations, butyrate cross-feeding between *F. plautii* and *C. catus* may support microbial diversity and metabolic cooperation.

Recent studies have reported the beneficial effects of *F. plautii* in murine models of myocardial ischemia / reperfusion injury, Th2 responses induced by antigens and the constitution of phlegm dampness (PDC) [11, 26, 37]. Although the underlying mechanisms remain unclear, our modeling supports the hypothesis that SCFA production, particularly butyrate, may contribute to these protective outcomes. These findings provide a rationale for exploring *F. plautii* as a candidate for probiotic development, with potential applications in modulating host immunity, barrier function, and microbial composition.

### Toward improved GEMs for gut microbiome research

Although *iFP655* achieved strong predictive performance for *F. plautii*, our reconstruction pipeline was less successful for other community members. This highlights ongoing challenges in GEM development for diverse and understudied gut microbes. Our results emphasize the importance of incorporating condition-specific information, such as medium constraints, thermodynamic feasibility, transcriptomic profiles, and biologically justified gap filling, to improve model accuracy.

As microbiome research increasingly integrates multi-omics data, high-quality GEMs will be essential for linking microbial metabolism to community dynamics and host outcomes. The workflow and insights presented in this study offer a roadmap for the generation of predictive models and highlight *F. plautii* as both a tractable modeling target and a functionally relevant member of the human gut ecosystem.

In conclusion, this study demonstrated that the curated model of *F. plautii, iFP655*, effectively predicts growth and butyrate production under defined intestinal conditions. By integrating thermodynamic constraints, transcriptomics data, and key biosynthetic pathways, particularly the lysine utilization route, the model offers improved accuracy over existing GEMs. Community simulations revealed that *F. plautii* plays a functional role in the production of short-chain fatty acids and amino acid metabolism, while supporting the syntrophic interactions shaped by the diet. Future research should experimentally validate these predictions, particularly under conditions mimicking diets low in fermentable carbohydrates but enriched in protein and lysine derivatives. Such approaches could confirm the potential probiotic roles of *F. plautii* in modulating gut health via SCFA production and lysine metabolism. However, the findings in this work highlight the ecological importance of *F. plautii* in gut microbial networks and strengthen its potential as a target for probiotic interventions. Moreover, this work underscores the need for condition-aware reconstruction frameworks that incorporate medium definition, thermodynamics, and omics integration to improve model fidelity for less-studied gut microbes.

## Methods

This study consisted of two major stages: (1) reconstruction and evaluation of a genome-scale model (GEM) for *F. plautii*, and (2) community modeling of *F. plautii* alongside other members of the gut microbiota within the Diet-based Minimal Microbiome (DbMM) community.

### Reconstruction and evaluation of the **F. plautii** GEM

#### In silico medium composition

The *in silico* medium was based on the DbMM formulation described previously [49]. This medium includes xylan, starch (potato), inulin (chicory), pectin (apple, and cellobiose as carbon sources. Additional components such as casitone, beef extract and peptone were approximated using the TSB composition, while amino acid uptake rates were set at 0.5 mmol/gDW/h. Where maximum flux data were unavailable, compound concentrations were used to estimate uptake limits following:

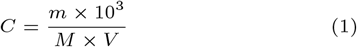

where *C* is concentration (mmol/L), *m* is mass (g), *M* is molecular weight (g/mol), and *V* is volume (L). The resulting DbMM composition is listed in the supplementary material and summarized in Table S1. AF medium data were taken from Weiss et al. [55], and M2 medium was estimated similarly.

#### Draft model reconstruction

The *F. plautii* genome (RefSeq: GCF 000242155.1) was used as input for CarveMe (v1.6.1) and Gapseq (v1.2) [29, 56]. Both reconstructions were gap-filled using DbMM medium. Manual refinement included: (i) correcting imbalanced reactions, (ii) removing duplicates, (iii) adding SCFA transport reactions, and (iv) constraining directionality using Gibbs free energy changes. The quality of the model was assessed with MEMOTE (v0.17.0) [28].

#### Thermodynamic constraint integration

Gibbs free energy changes (Δ_*r*_*G*^′^) were calculated under gut-like conditions (pH 6, 310K, ionic strength 0.25 M) using eQuilibrator v3.0 [3]. Reactions with Δ_*r*_*G*^′^ between *−*30 and +30 kJ/mol were set as reversible; reactions outside this range were constrained accordingly [5, 47].

#### Gap-filling with CHESHIRE

CHESHIRE [8], a deep learning-based gap-filling tool, was applied with DbMM medium as input. Reactions in the top 25, 50, 100, and 200 were iteratively added and directionality adjusted based on Δ_*r*_*G*^′^. The models were screened to exclude reactions that formed energy-generating cycles or caused unrealistic flux inflation.

#### Model evaluation

Parsimonious Flux Balance Analysis (pFBA) was performed using COBRApy (v0.29.0) [12] and GLPK. The cellobiose exchange reaction was constrained to 0 mmol/gDW/h based on experimental findings [48]. Simulations were performed in DbMM, AF, and M2 media. The model best aligned with the experimental growth and SCFA secretion data was selected for further analysis. Flux Variability Analysis (FVA) was used to estimate the permissible flux ranges that achieve at least 90% of optimal growth.Gene and Reaction Essentiality Analysis was conducted using `single_gene_deletion` and `single_reaction_deletion` functions in COBRApy to identify essential components.

#### Transcriptomic data integration

Transcriptomic data from a 10-species community of DbMM [48] were filtered to retain *F. plautii*-specific genes. CPM normalization was applied and the top 30 percentile of genes was integrated using the GIMME algorithm through the Troppo package [14]. Low-flux reactions (non-essential, non-spontaneous, gene-associated) were knocked out. The integration improved pathway contextualization and improved flux predictions.

### DbMM community modeling

#### Community member GEMs reconstruction

The GEMs for other DbMM species were reconstructed using the optimized pipeline that produced *iFP655*. Genomes were retrieved from NCBI (RefSeq IDs in the supplementary material, Table S2). The predicted growth rates were compared against experimental data [48].

#### Three-species community modeling

A three-member community of *F. plautii, B. ovatus*, and *C. catus* was simulated under DbMM conditions using PyCoMo (v0.2.6) [39]. Species abundances (0.15, 0.59, 0.26 respectively) were derived from prior 10-member community data [48]. Oxygen exchange was blocked, and the community biomass function was maximized. FBA and FVA were performed, and potential cross-feeding interactions were visualized in Cytoscape using ScyNet [38].

## Competing interests

No competing interest is declared.

## CRediT authorship contribution statement

**William T. Scott, Jr**.: Writing – review & editing, Writing – original draft, Visualization, Validation, Supervision, Methodology, Investigation, Formal analysis, Conceptualization.

**Enden Dea Nataya**: Writing – original draft, Visualization, Validation, Methodology, Investigation, Formal analysis, Conceptualization.

**Clara Belzer**: Writing – review & editing, Conceptualization.

**Peter J. Schaap**: Writing – review & editing, Project administration, Funding acquisition. Conceptualization.

## Data and code availability statement

All scripts, final models, and supplementary data used or generated in this study are available at the following GitLab repository: https://git.wur.nl/ssb/student-projects/enden_dea_nataya_thesis_msc. Furthermore, the GEMs, as well as the accompanying MEMOTE [28] and FROG Analysis [42] reports, can be found in a public repository on BioModels: https://www.ebi.ac.uk/biomodels/.

## Acknowledgments

W.T.S.J. and P.J.S. acknowledge the Dutch Research Council (NWO) and Wageningen University & Research for their financial contribution to UNLOCK (NWO: 184.035.007).

## Notes

### Competing Interest Statement

The authors have declared no competing interest.

### Summary of Updates

There was a typo in the abstract, and one sentence in the discussion needed further clarity in explaining the reactions removed.

