## Supplementary Material for "Metabolic modeling unveils potential probiotic roles of *Flavonifractor plautii* in reshaping the Western gut microbiota landscape" (from another version of this preprint: version 1, 2025-04-21)

### A.1 Appendix 1: The DbMM medium composition

***Table S1*** *The DbMM medium composition*

| medium | name | bigg | modelseed | maxflux |
|---|---|---|---|---|
| DbMM | Potassium (K) | k | cpd00205 | 9.122323 |
| DbMM | H2PO4 | pi | cpd00009 | 7.674393 |
| DbMM | Sodium (Na) | na1 | cpd00971 | 91.57618 |
| DbMM | NH4 | nh4 | cpd00013 | 6.103 |
| DbMM | CL- | cl | cpd00099 | 14.574 |
| DbMM | Magnesium (Mg) | mg2 | cpd00254 | 0.716 |
| DbMM | HCO3- | hco3 | cpd00242 | 47.6 |
| DbMM | H+ | h | cpd00067 | 0.05 |
| DbMM | Manganese (Mn) | mn2 | cpd00030 | 0.551262 |
| DbMM | FE2+ | fe2 | cpd10516 | 0.6375 |
| DbMM | Co2+ | cobalt2 | cpd00011 | 0.0005 |
| DbMM | Ni2+ | ni2 | cpd00244 | 0.0001 |
| DbMM | Zn2+ | zn2 | cpd00034 | 3.6405 |
| DbMM | Cu2+ | cu2 | cpd00058 | 0.1147 |
| DbMM | OH- | h2o | cpd00001 | 100 |
| DbMM | SeO32- | slnt | cpd03387 | 0.0001 |
| DbMM | WO4- | tungs | cpd15574 | 0.0001 |
| DbMM | MoO4- | mobd | cpd11574 | 0.0001 |
| DbMM | Cysteine- | cys__L | cpd00084 | 0.5 |
| DbMM | Haemin | hemeA | cpd04145 | 7.67E-05 |
| DbMM | Vit K | phllqne | cpd01401 | 0.001 |
| DbMM | pectin | pectin | cpd11601 | 0.0067 |
| DbMM | Inulin |  | cpd11602 | 0.2 |
| DbMM | Xylan | xylan4 | cpd11732 | 0.006 |
| DbMM | Cellobiose | cellb | cpd11746 | 2.92 |
| DbMM | Starch | starch | cpd11657 | 0.0228 |
| DbMM | Biotin | btn | cpd00104 | 0.00082 |
| DbMM | Nicotinamide | ncam | cpd00133 | 0.01638 |
| DbMM | p-Aminobenzoic acid | anth | cpd00093 | 0.00729 |
| DbMM | Thiamine | thm | cpd00305 | 0.06654 |
| DbMM | Pantothenic acid | pnto__R | cpd00644 | 0.23256 |
| DbMM | Pyridoxamine | pydam | cpd00419 | 0.02955 |
| DbMM | Cyanocobalamin (B12) | b12 | cpd01826 | 0.00074 |
| DbMM | Riboflavin | ribflv | cpd00220 | 0.18866 |
| DbMM | CH3COO- | ac | cpd00029 | 30 |
| DbMM | Aspartic acid | asp__L | cpd00041 | 0.5 |
| DbMM | Glutamic acid | glu__L | cpd00023 | 0.5 |
| DbMM | Asparagine | asn__L | cpd00132 | 0.5 |
| DbMM | Glutamine | gln__L | cpd00053 | 0.5 |

| | | | | |
|---|---|---|---|---|
| DbMM | Alanine | ala__L | cpd00035 | 0.5 |
| DbMM | Arginine | arg__L | cpd00051 | 0.5 |
| DbMM | Glycine | gly | cpd00033 | 0.5 |
| DbMM | Histidine | his__L | cpd00119 | 0.5 |
| DbMM | Isoleucine | ile__L | cpd00322 | 0.5 |
| DbMM | Leucine | leu__L | cpd00107 | 0.5 |
| DbMM | L-Lysine | lys__L | cpd00039 | 0.5 |
| DbMM | L-Methionine | met__L | cpd00060 | 0.5 |
| DbMM | Phenylalanine | phe__L | cpd00066 | 0.5 |
| DbMM | Proline | pro__L | cpd00129 | 0.5 |
| DbMM | L-Serine | ser__L | cpd00054 | 0.5 |
| DbMM | Threonine | thr__L | cpd00161 | 0.5 |
| DbMM | Tyrosine | tyr__L | cpd00069 | 0.5 |
| DbMM | Valine | val__L | cpd00156 | 0.5 |
| DbMM | Calcium (Ca) | ca2 | cpd00063 | 0.30568 |
| DbMM | Chromium (Cr) | cro2 | cpd12859 | 0.00731 |
| DbMM | Cobalt (Co) | cobalt2 | cpd00149 | 0.0102 |
| DbMM | Molybdenum (Mo) | mobd | cpd00131 | 0.000625 |
| DbMM | Selenium (Se) | slnt | cpd01079 | 0.0076 |
| DbMM | Nicotinic acid (B3) | nac | cpd00218 | 19.51254 |
| DbMM | Pyridoxine (B6) | pydx5p | cpd00016 | 0.065515 |
| DbMM | Folic acid (B9) | fol | cpd00393 | 0.000136 |
| DbMM | L-Tryptophan | trp__L | cpd00065 | 0.5 |
| DbMM | Iron (Fe) | fe3 | cpd10515 | 0.844608 |
| DbMM | Creatinine | crtn | cpd00585 | 2.482 |
| DbMM | Vitamin B12 | adocbl | cpd00166 | 0.00077 |
| DbMM | Taurine | taur | cpd00210 | 0.0511 |
| DbMM | Citrulline | citr__L | cpd00274 | 0.0297 |
| DbMM | Carnosine | carn | cpd15836 | 0.327 |
| DbMM | Anserine | ans | cpd00926 | 0.0624 |
| DbMM | Creatine | creat | cpd00250 | 0.732 |
| DbMM | Hypoxanthine | hxan | cpd00226 | 0.279 |
| DbMM | Inosine | ins | cpd00246 | 0.0522 |
| DbMM | Lactic acid | lac__L | cpd00159 | 3.243 |
| DbMM | Glycolic acid | glyclt | cpd00139 | 0.258 |
| DbMM | Succinic acid | succ | cpd00036 | 0.213 |
| DbMM | Carnitine | crn | cpd00266 | 0.409 |
| DbMM | Urea | urea | cpd00073 | 0.0366 |
| DbMM | Sulphate | so4 | cpd00048 | 0.001 |
| DbMM | Benzoate | bz | cpd00153 | 0.01 |

### A.2 Appendix 2: The DbMM community members

***Table S2*** *The DbMM community members and their NCBI genome accession number*

| No. | Species | NCBI Genome Accession |
|---|---|---|
| 1 | *Agathobacter rectalis* | GCF_000020605.1 |

| | | |
|---|---|---|
| 2 | *Anaerobutyricum soehngenii* | GCF_009697165.1 |
| 3 | *Bacteroides ovatus* | GCF_001314995.1 |
| 4 | *Bacteroides xylanisolvens* | GCF_000162155.1 |
| 5 | *Coprococcus catus* | GCF_019734885.1 |
| 6 | *Eubacterium siraeum* | GCF_000382085.1 |
| 7 | *Faecalibacterium prausnitzii* | GCF_000154385.1 |
| 8 | *Flavonifractor plautii* | GCF_000242155.1 |
| 9 | *Roseburia intestinalis* | GCF_000156535.1 |
| 10 | *Subdoligranulum variabile* | GCF_000157955.1 |

### A.3 Appendix 3: Essential Genes in *i*FP655

***Table S3*** *Essential genes in iFP655*

| No | GPR | Gene id | General Subsytem | ePath database |
|---|---|---|---|---|
| 1 | WP_007488992_1 | purB | Purine and pyrimidine metabolism | Yes |
| 2 | WP_009258108_1 | pgk | Carbohydrates metabolism | Yes |
| 3 | WP_007495936_1 | prsA | Carbohydrates metabolism | Yes |
| 4 | WP_009256962_1 | ddl | Amino acids metabolism | Yes |
| 5 | WP_007488232_1 | metF | Carbohydrates metabolism | Yes |
| 6 | WP_009256893_1 | | Vitamins and cofactors biosynthesis | No |
| 7 | WP_140402426_1 | pncB | Vitamins and cofactors biosynthesis | Yes |
| 8 | WP_007489024_1 | gmk | Purine and pyrimidine metabolism | Yes |
| 9 | WP_009260498_1 | xdhB | Purine and pyrimidine metabolism | Yes |
| 10 | WP_007489118_1 | lysC | Amino acids metabolism | Yes |
| 11 | WP_007489388_1 | | Vitamins and cofactors biosynthesis | No |
| 12 | WP_009258949_1 | dapF | Amino acids metabolism | Yes |
| 13 | WP_007495532_1 | dxr | Vitamins and cofactors biosynthesis | Yes |
| 14 | WP_009260793_1 | guaA | Purine and pyrimidine metabolism | Yes |
| 15 | WP_009260380_1 | glmS | Amino acids metabolism | Yes |
| 16 | WP_110503348_1 | | Carbohydrates, peptidoglycans, and teichoic acids | No |
| 17 | WP_009258099_1 | nadE | Vitamins and cofactors biosynthesis | Yes |
| 18 | WP_009259022_1 | | Amino acids metabolism | No |
| 19 | WP_009258109_1 | tpiA | Carbohydrates metabolism | Yes |
| 20 | WP_007495529_1 | ispG | Vitamins and cofactors biosynthesis | Yes |
| 21 | WP_009257515_1 | pyrE | Purine and pyrimidine metabolism | Yes |
| 22 | WP_007493707_1 | GXM20_RS02945 | Transport reactions | No |
| 23 | WP_009259232_1 | nadD | Vitamins and cofactors biosynthesis | Yes |
| 24 | WP_009259163_1 | | Transport reactions | No |
| 25 | WP_007494624_1 | adk | Purine and pyrimidine metabolism | Yes |
| 26 | WP_009256823_1 | | Vitamins and cofactors biosynthesis | No |
| 27 | WP_009260813_1 | fbp3 | Carbohydrates metabolism | Yes |
| 28 | WP_009256961_1 | murF | Amino acids metabolism | Yes |
| 29 | WP_007495705_1 | coaD | Vitamins and cofactors biosynthesis | Yes |
| 30 | WP_021631186_1 | dxr | Vitamins and cofactors biosynthesis | Yes |
| 31 | WP_009257520_1 | murC | Carbohydrates, peptidoglycans and teichoic acids | Yes |
| 32 | WP_009260571_1 | | Carbohydrates metabolism | Yes |

| | | | | |
|---|---|---|---|---|
| 33 | WP_007495534_1 | | Vitamins and cofactors biosynthesis | No |
| 34 | WP_007489380_1 | asd | Amino acids metabolism | Yes |
| 35 | WP_007489383_1 | dapB | Amino acids metabolism | Yes |
| 36 | WP_009258111_1 | gpml | Carbohydrates metabolism | Yes |
| 37 | WP_009257212_1 | | Carbohydrates metabolism | No |
| 38 | WP_009260499_1 | xdhA | Purine and pyrimidine metabolism | Yes |
| 39 | WP_009257907_1 | | Amino acids metabolism | No |
| 40 | WP_009256752_1 | pyrC | Purine and pyrimidine metabolism | Yes |
| 41 | WP_009260497_1 | xdhC | Purine and pyrimidine metabolism | No |
| 42 | WP_021632369_1 | ispF | Vitamins and cofactors biosynthesis | Yes |
| 43 | WP_009260538_1 | coaBC | Vitamins and cofactors biosynthesis | Yes |
| 44 | WP_009257485_1 | ribF | Vitamins and cofactors biosynthesis | Yes |
| 45 | WP_009256844_1 | glmM | Amino sugar and nucleotide sugar metabolism | Yes |
| 46 | WP_007493348_1 | crt | Amino acids metabolism | Yes |
| 47 | WP_009258317_1 | thyA | Purine and pyrimidine metabolism | Yes |
| 48 | WP_009259625_1 | pyrG | Purine and pyrimidine metabolism | Yes |
| 49 | WP_009257631_1 | purA | Purine and pyrimidine metabolism | Yes |
| 50 | WP_007489711_1 | ispE | Vitamins and cofactors biosynthesis | Yes |
| 51 | WP_007495720_1 | mraY | Carbohydrates, peptidoglycans and teichoic acids | Yes |
| 52 | WP_138307007_1 | murD | Amino acids metabolism | Yes |
| 53 | WP_009261245_1 | gapA | Vitamins and cofactors biosynthesis | Yes |
| 54 | WP_009256753_1 | pyrF | Purine and pyrimidine metabolism | Yes |
| 55 | WP_009260807_1 | tktA | Carbohydrates metabolism | Yes |
| 56 | WP_049893253_1 | ispD | Vitamins and cofactors biosynthesis | Yes |

### A.4 Oxygen-dependent reactions that were essential for e_siraeum, s_variabile, and b_ovatus model

***Table S4*** *Essential oxygen-dependent reactions in e_siraeum, s_variabile, and b_ovatus model*

| Model | Reaction id |
|---|---|
| e_siraeum | ALOX |
| e_siraeum | AHMMPS_1 |
| e_siraeum | FAS181 |
| s_variabile | CYTBD2pp |
| s_variabile | INOSTO |
| b_ovatus | AHMMPS_1 |
| b_ovatus | PDX5POi |
| b_ovatus | CAT |
